# Comparative analysis of intragroup intermale relationships: A study of wild bonobos (*Pan paniscus*) in Wamba, Democratic Republic of Congo and chimpanzees (*Pan troglodytes*) in Kalinzu Forest Reserve, Uganda

**DOI:** 10.1101/2023.09.10.557020

**Authors:** Shohei Shibata, Takeshi Furuichi

**Affiliations:** Wildlife Research Center, Kyoto University, Inuyama, Aichi 484-8506, Japan

**Keywords:** *Pan paniscus*, *Pan troglodytes*, aggression, fission–fusion dynamics, party size, dominance rank, Wamba, Kalinzu Forest Reserve

## Abstract

Although chimpanzees (*Pan troglodytes*) and bonobos (*Pan paniscus*) share a multi- male/multi-female societal organization and form male-philopatric groups, disparities in terms of male aggression and stability of temporary parties are thought to exist among them. However, existing research in bonobos has mainly focused on the high social status, prolonged receptivity, and characteristic sexual behaviors of females, leaving the behaviors of males understudied. Moreover, prior comparative studies on *Pan* suffer from methodological inconsistencies. This study addresses these gaps by employing a uniform observation method to explore party attendance and aggressive interactions among male bonobos in Wamba and male chimpanzees in Kalinzu. Unlike male chimpanzees that exhibit dispersion in the absence of receptive females in the group, male bonobos showed a lesser degree of such dispersion. Although the overall frequency of aggressive interactions per observation unit did not significantly differ between the two species, the nature of these interactions varied. Notably, severe aggressive behaviors such as physical confrontations among adult males were absent in bonobos, with most aggression occurring between the sons of the two highest-ranking females. Additionally, in bonobos, females actively engaged in polyadic aggressive behavior as aggressors, while all instances of coalitionary aggression in chimpanzees originated from male aggressors. These findings underscore the substantial impact of female behaviors on the observed distinctions in male aggressive interactions between the two species.

## Introduction

Factors concerning intermale mating competition primarily influence the reproductive strategies and social relationships of primate males. These factors include the number of receptive females, mating seasonality, operational sex ratio, dominance rank among males in a group, and relationships with other groups or out-group males. Among male chimpanzees (*Pan troglodytes*), the dominance hierarchy is relatively linear, and aggressive behaviors from dominant individuals to subordinate individuals are commonly observed in many dyads (Nishida and Hosaka 1996; Newton-Fisher 2004). The aggression among males seems to be related to male reproductive strategies because the presence of receptive females increases male aggression (Muller and Wrangham 2004). The aggressive behaviors are sometimes very violent and may result in the killing of adult males of the same group (Fawcett and Muhumuza 2000; Watts 2004; Kaburu et al. 2013; Wilson et al. 2014). On the other hand, some male members of a chimpanzee group form coalitions, which may help them monopolize mating opportunities against other male members (Watts 1998; Duffy et al. 2007; Mitani 2009; Schülke et al. 2010).

Bonobos (*Pan paniscus*) form male-philopatric groups that are very similar to those of chimpanzees (Kano 1982; Gerloff et al. 1999; Hashimoto et al. 2008; Ishizuka et al. 2019). Therefore, we can expect to see similar behaviors in males, including intense aggression against other males in the group, an increase in the aggressive interactions in the presence of receptive females, coalitions among males for the monopolization of receptive females. However, these behaviors are seldom observed or are observed with less intense aggression among males within a group (Kano 1992; Furuichi and Ihobe 1994; Furuichi 1997; Surbeck et al. 2012; Samuni et al. 2022), and it raises the question of why bonobos and chimpanzees show substantial differences in their aggression and behavior patterns. Moreover, a significant question arises regarding the prevalence of reproductive skew toward high-ranking males in bonobos compared to chimpanzees, even though the intermale relationships in bonobos appear to be less despotic (Surbeck et al. 2017b; Ishizuka et al. 2018; Surbeck et al. 2019; Yokoyama and Furuichi 2022; Ishizuka 2023).

While answering this question requires observations of males from both species, studies focusing on behavioral interactions among male bonobos have been relatively scarce (Kuroda 1980; Ihobe 1992; Furuichi and Ihobe 1994; Surbeck et al. 2017a; Cheng et al. 2022). This scarcity is likely attributed to the less conspicuous nature of male bonobos behaviors compared to those of chimpanzees. Additionally, the heightened conspicuousness of female behavioral traits in bonobos may contribute to the focus on the latter (Parish 1994; White and Wood 2007; Furuichi 2011, 2023). Furthermore, most previous studies comparing the behavior of wild chimpanzees and bonobos did not rely on data obtained through uniform observation methods. Therefore, to clarify whether there are significant differences in within-group male relationships between chimpanzees and bonobos, we conducted studies using the same observation methods under natural conditions of male bonobos in Wamba, Democratic Republic of the Congo and male chimpanzees in Kalinzu Forest Reserve, Uganda.

In contrast to chimpanzees, bonobos are not male dominant but rather show male-female co dominance (Furuichi 1997; Stevens et al. 2007). The patterns of association and behavior of male bonobos have been considered to be strongly affected by prolonged sexual receptivity of females and the constant presence of multiple receptive females in parties (Furuichi and Hashimoto 2002; Hashimoto et al. 2022), moderation of mating competition by the presence of multiple receptive females (Kano 1992; Furuichi and Hashimoto 2002; Furuichi 2023), long-lasting mother–son relationships (Furuichi 1989; Kano 1992; Furuichi 2011), and strong support from dominant mothers in mating competition and acquisition of alpha status (Surbeck et al. 2010; Furuichi 2011; Ishizuka et al. 2018; Furuichi 2019; Surbeck et al. 2019; Shibata and Furuichi 2023). Based on these previous reports, we hypothesized that in bonobos, unlike chimpanzees, the prolonged sexual receptivity of females and the support from high-ranking mothers strongly influence agonistic relationships among males so that the importance of intermale aggressive interactions is relatively lower, and the frequency and intensity of male aggression are reduced. In this study, we tested four predictions derived from our hypothesis.

1. Male bonobos are more likely to find a higher number of receptive females in parties to which they attend than do male chimpanzees.
2. In bonobos, instances of aggressive interactions among males are less frequent or involve less physical contact than in chimpanzees.
3. Aggressive interactions between males over access to receptive females are less frequent in bonobos than in chimpanzees.
4. In bonobos, aggressive interactions between males more commonly involve their mothers as coalition partners compared to the patterns observed in chimpanzees.

## Methods

### Study sites and subjects

#### Bonobos in Wamba

Field assistants and author S.S. observed wild bonobos of the E1 group in Wamba in the northern sector of the Luo Scientific Reserve in DR Congo, where long- term research has been conducted since 1973 (Kano 1992; Furuichi et al. 2012). The bonobos were observed for 37 days from September 19 to December 29, 2019. All the individuals in the E1 group were identified and habituated before the beginning of this study. At the time of this research, the group comprised approximately 40 individuals, including eight adult males aged over 14 years, three adolescent males aged 8–14 years, 13 adult females, a few immigrant adolescent females, and many juveniles and infants. The age classes of subject males were categorized according to Hashimoto (1997). The study subjects were eight adult and three adolescent males. The individual names, dominance rank, estimated birth year, and their mothers are shown in Table 1. The dominance rank of males was calculated using David’s score (David 1987) based on dyadic aggressive interactions (see below for the definition of aggressive interactions). The individuals were divided into one of the following four categories, namely the alpha male (1), high ranking (2–3), middle ranking (4–7), and low ranking (8–11).

**Table 1.**
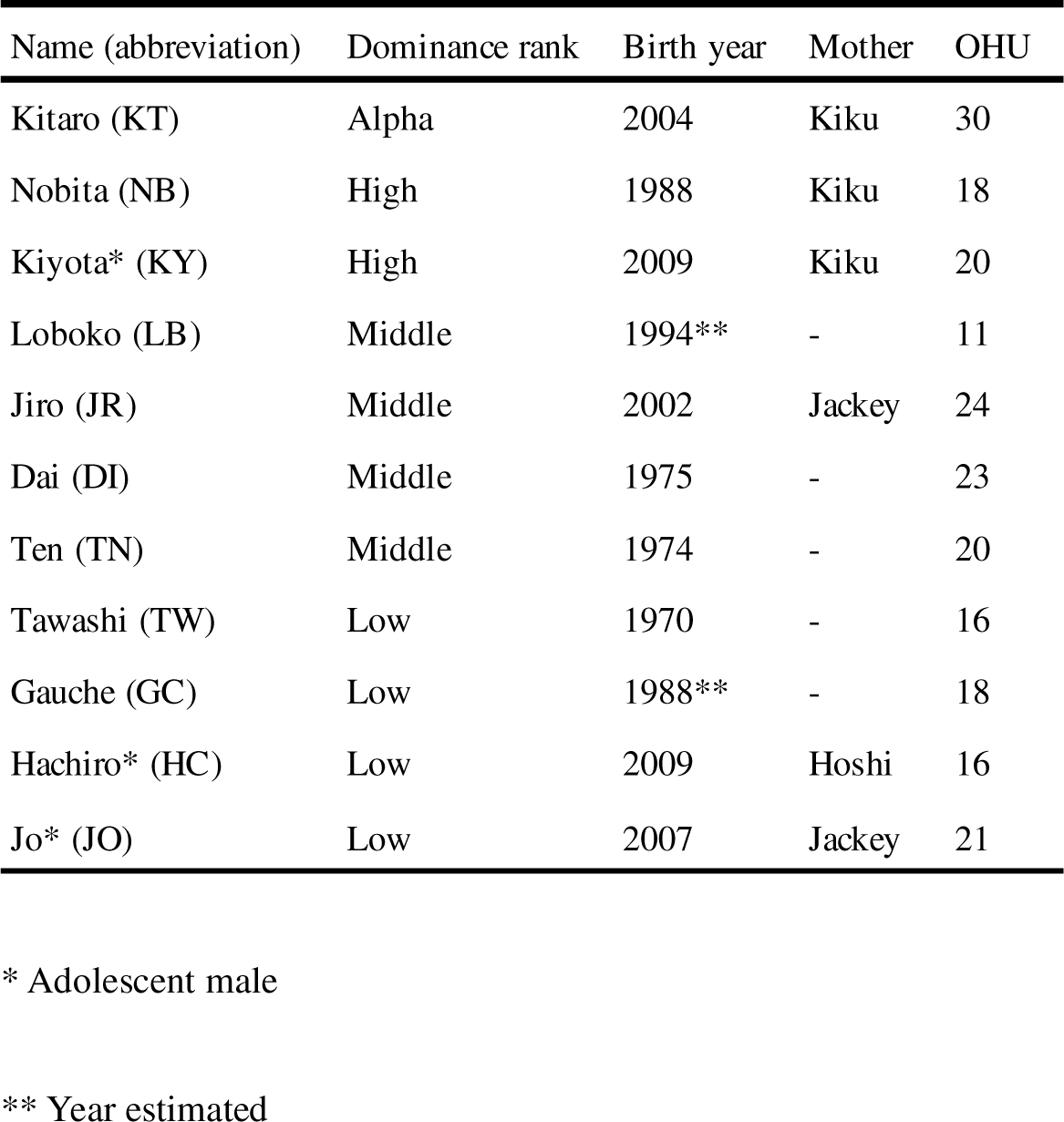
Subject males, dominance rank, birth year, and the number of observation hour units (OHU) of bonobos of the E1 group.

| Name (abbreviation) | Dominance rank | Birth year | Mother | OHU |
| --- | --- | --- | --- | --- |
| Kitaro (KT) | Alpha | 2004 | Kiku | 30 |
| Nobita (NB) | High | 1988 | Kiku | 18 |
| Kiyota* (KY) | High | 2009 | Kiku | 20 |
| Loboko (LB) | Middle | 1994** | - | 11 |
| Jiro (JR) | Middle | 2002 | Jackey | 24 |
| Dai (DI) | Middle | 1975 | - | 23 |
| Ten (TN) | Middle | 1974 | - | 20 |
| Tawashi (TW) | Low | 1970 | - | 16 |
| Gauche (GC) | Low | 1988** | - | 18 |
| Hachiro* (HC) | Low | 2009 | Hoshi | 16 |
| Jo* (JO) | Low | 2007 | Jackey | 21 |

\* Adolescent male
\*\* Year estimated

#### Chimpanzees in Kalinzu

Field assistants and author S.S. observed the wild chimpanzees of the M group in the Kalinzu Forest Reserve, Uganda, where long-term research has been conducted since 1992 (Hashimoto 1995; Hashimoto and Furuichi 2006). The chimpanzees were observed for 112 days from February 3 to April 18, from June 23 to September 1, 2018, and from March 11 to May 24 2019. During this study, the group consisted of approximately 100 individuals, including 15 adult males aged over 15 years (Goodall 1986), 29 adult females, a few adolescent males and females, and many juveniles and infants. All adult males of the M group were identified and habituated by the beginning of the study period. The study subjects of the current study comprised 10 adult males. The individual names, dominance ranks, and estimated birth years of each adult male are shown in Table 2. The dominance rank of males was calculated using David’s score (David 1987) based on dyadic aggressive interactions. The individuals were divided into one of the following four categories, namely the alpha male (1), high ranking (2–4), middle ranking (5–7), and low ranking (8–10) (Table 2).

**Table 2.** Subject males, dominance rank, birth year, and the number of observation hour units (OHU) of chimpanzees of the M group.

| Name (abbreviation) | Dominance rank | Birth year | OHU |
| --- | --- | --- | --- |
| Goku (GK) | Alpha | 1993* | 90 |
| Ponta (PO) | High | 1995* | 53 |
| Ichiro (IC) | High | 1980s* | 76 |
| Buru (BR) | High | 1970s* | 65 |
| Prince (PR) | Middle | 1997* | 59 |
| Taiki (TK) | Middle | 1999 | 77 |
| Deo (DO) | Middle | 1970s* | 62 |
| Pietan (PT) | Low | 2001 | 65 |
| Black (BL) | Low | 1998* | 72 |
| Jo (JO) | Low | 2000* | 58 |

\* Year estimated

#### Data collection

Bonobos in Wamba were followed from approximately 0600 h from their previous night’s sleep site. Chimpanzees in Kalinzu were followed daily from approximately 0700 h. We conducted observations using the same methods for both bonobos and chimpanzees. We used the focal sampling method (Altmann 1974), in which a focal animal was followed for as long as possible. We followed the first male we found in the morning unless we had followed that individual on the previous day. When we lost sight of the focal individual, we continued to observe if the individual was found again within 30 min. However, if we could not find the individual for 30 min, we stopped the focal observation for that day.

#### Party composition and observation unit

Every 60 min during the focal following, we recorded the party composition that the focal animal belonged to using the 1-h party method (Hashimoto et al. 2001; Mulavwa et al. 2008). We recorded individuals within the visible range at the beginning of each hour and included new individuals that joined the party until the end of the hour. Each hour-long observation was considered a unit for analysis and referred to as one observational hour unit (OHU) when focal animals were observed for > 30 min within that hour. To exclude the potential for overestimating data on behaviors that were easily observed, we excluded all observation data for days with < 3 OHUs (11 OHUs for bonobos and 22 OHUs for chimpanzees). In total, 217 OHUs were analyzed for all the subject bonobos with a mean of 19.7 ± 5.0 SD OHUs (range: 11–30) for each individual (Table 1). The mean duration of OHU was 53.5 min (range: 30–60). For all the chimpanzee subjects, 677 OHUs were analyzed with a mean of 67.7 ± 11.1 SD OHUs (range: 53–90) for each individual (Table 2). The mean duration of OHU was 52.1 min (range: 30–60).

#### Aggressive interactions

We recorded all the observed intragroup aggressive interactions in each OHU using *ad libitum* sampling, including cases where the focal animal was not involved. We defined “aggressive interactions” as involving at least one aggressive behavior toward one or more individuals: aggressive physical contact such as kicking, beating, and biting; chasing; charging; directed aggressive displaying (including branch dragging and provocative behaviors for bonobos); non-vocal threatening. Undirected aggressive displays, namely jumping in chimpanzees and branch dragging in bonobos, were only recorded as aggressive interactions when other individuals responded by screaming or fleeing after that behavior. Interactions involving only pant-grunts in chimpanzees were not recorded as aggressive interactions because the frequency of such interactions was very low in chimpanzees in Kalinzu, and bonobos did not exhibit corresponding behaviors. Although we could not record some aggressive interactions due to bad visibility in the bush or on the high canopy, all recorded aggressive interactions that involved subject males were included in the analyses. The visibility was not substantially different between the two study sites; therefore, it did not bias comparisons between bonobos and chimpanzees in this study.

#### Presence of females with maximum swelling and comparison of party attendance by adult males

The swelling status of the sexual skin of each female in each party was recorded as one of the following two categories: non-swelling and maximum swelling (firmness 3 defined by (Furuichi 1987), hereinafter MS).

We defined all OHUs on days when we observed at least one female with MS as “OHU under the presence of females with MS.” In bonobos, females resume receptivity in the early stage of postpartum infertility; therefore, usually multiple females show MS and sexual receptivity (Kano 1992; Furuichi and Hashimoto 2002; Hashimoto et al. 2022). In the current study, bonobo females with MS were observed in 35 of 37 (97%) observation days, and 197 of 217 (90.8%) OHUs were defined as “OHU under the presence of females with MS.” Therefore, we included all 217 OHUs in bonobos as a single dataset in the analysis of the party attendance by adult males. In chimpanzees, females with MS were observed in 62 of 112 (55.4%) observation days; 376 out of 677 (55.5%) OHUs were defined as “OHU under the presence of females with MS,” and the remaining 301 OHUs were defined as “OHU under the absence of females with MS.” We compared the number of adult males observed in a party in three different group conditions, namely bonobos, chimpanzees under the presence of females with MS, and chimpanzees under the absence of females with MS (Fig. 1). To determine whether the datasets of the three conditions followed the same distribution, we first normalized each dataset and then conducted the Kolmogorov–Smirnov two- sample tests for three combinations by using the “ks.test()” function. We used R (version 4.2.2; R Foundation for Statistical Computing, Vienna, Austria, http://www.r-project.org [accessed on 13 December 2022]) for all statistical analyses.

**Figure 1.**
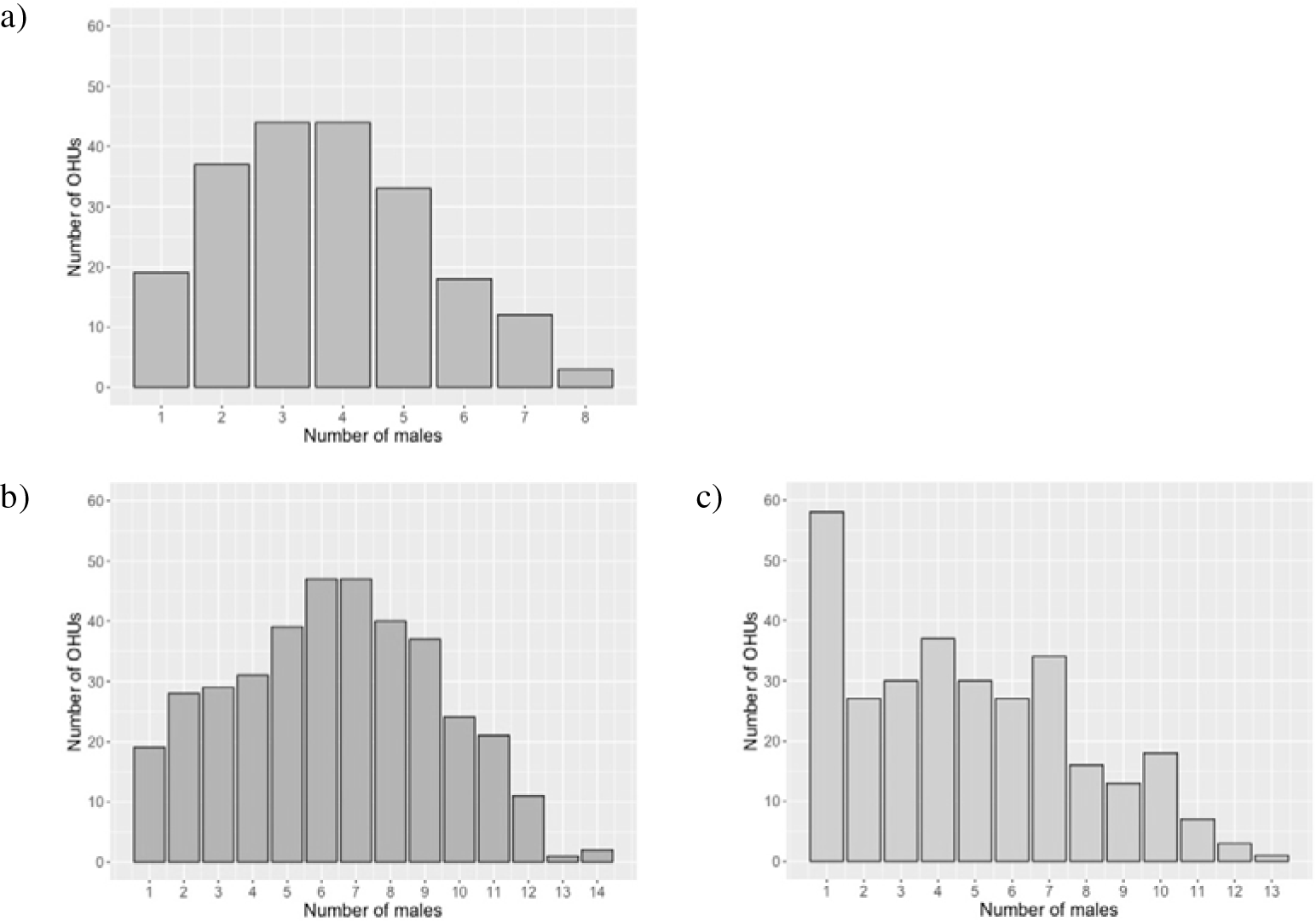
The number of OHUs in which the party included the respective number of adult males. a) Adult male bonobos (N = 217). b) Adult male chimpanzees under the presence of females with MS (N = 376). c) Adult male chimpanzees under the absence of females with MS (N = 301).

#### Intragroup aggressive interactions among males

We recorded 92 aggressive interactions among male bonobos. Among these, 76 cases were dyadic aggressive interactions between males, comprising 43 cases in 12 dyads between adult males and 33 cases in 10 dyads between adult and adolescent males. The remaining 16 cases were aggression toward males from multiple individuals (including at least one male as an aggressor), comprising 14 cases toward adult males and two cases toward adolescent males. For chimpanzees, we recorded 201 aggressive interactions among adult males, including 177 dyadic aggressive interactions in 65 dyads and 24 polyadic interactions. Polyadic interactions include both cases of coalitions and cases when there were multiple victims. We divided aggressive interactions into two conditions for chimpanzees based on the presence/absence of females with MS in the group. The aggressive interactions were divided into four categories according to their intensity, namely aggressive physical contact, chasing, directed aggressive display, and others including non-vocal threatening. All the aggressive interactions among all the subject bonobos (including adolescents) were summarized in the aggressive interaction matrix.

We calculated the frequencies of aggressive interactions in two different ways. First, we calculated the frequency of overall aggressive interactions per OHU by dividing the number of overall aggressive interactions by the number of OHUs. Secondly, we calculated the frequencies of aggressive interactions per male in an OHU by dividing the total number of males involved in aggressive interactions (including both aggressors and receivers) by the total number of males observed during each OHU. To examine the difference in the frequency of aggressive interactions between each group/condition, we conducted a chi-square test with a Bonferroni-corrected *post hoc* test using the “pairwise.prop.test()” function.

## Results

### Availability of receptive females and grouping patterns of adult males

A large difference between bonobos and chimpanzees in the availability of receptive females with MS for males was observed. In bonobos, 1.22 ± 1.15 (range 0–4) females with MS were observed in each OHU. On the other hand, in chimpanzees, 0.47 ± 0.68 (range 0–3) females with MS were observed in each OHU.

Figure 1 shows the number of adult males observed in each OHU. In chimpanzees, the number of males in parties differed considerably between days when females with MS were observed (Fig. 1b) and days when they were not observed (Fig. 1c). In the absence of females with MS, the modal value of the number of males was one, indicating that the males were highly dispersed. For bonobos, we drew only one pooled figure because females with MS were absent only in two days (Fig. 1a). The tendency of a high degree of dispersion was not found in bonobos. The party size distribution differed significantly in all three pairs (Kolmogorov–Smirnov two sample tests, chimpanzees under the absence of females with MS vs. chimpanzees under the presence of them, D = 0.125, *p* = 0.011; bonobos vs. chimpanzees under the absence of females with MS, D = 0.163, *p* = 0.003; bonobos vs. chimpanzees under the presence of females with MS, D = 0.145, *p* = 0.006). However, the shape of the distribution for male bonobos and the one for male chipanzees under the presence of females with MS were similar with modal number of males of three to four (37.5–40.0% of adult male members) for bonobos and six to seven (40.0–46.7%) for chimpanzees, suggesting that both male bonobos and chimpanzees show similar tendency of party attendance when females with MS are present in the group.

### Aggressive interactions among male bonobos and chimpanzees

Table 3 shows the number of dyadic aggressive interactions expressed and received in each dyad among 11 adult and adolescent male bonobos. We included those involving adolescent males for bonobos because those aggressive interactions played substantial roles in the dominance struggles between mother-son groups. Dyadic aggressive interactions were observed in 22 out of 55 dyads. Of the 76 dyadic aggressive interactions, 37 cases were directed at two middle and low-ranking males, Jiro and Jo, and 30 out of the 37 were from one of the three high-ranking males (Kitaro, Nobita, and Kiyota). In the 16 cases of polyadic aggressive interactions involving more than two individuals (including 14 cases among adult males and two cases involving adolescent males), the aggressee was a single male in all these cases and 15 cases were directed at Jiro or Jo. Kitaro, Nobita, and Kiyota are the sons of the alpha female Kiku, and Jiro and Jo are the sons of the second-ranking female Jacky (Table 1). During this observation period, the sons of Jacky were challenging the high status of the sons of Kiku by provoking them, and aggressive interactions among them frequently involved their mothers and brothers (Shibata, personal observation).

**Table 3.** Aggressive interaction matrix with numbers of aggression expressed and received among 11 adult and adolescent male bonobos of the E1 group.

|  |  | Receiver |  |  |  |  |  |  |  |  |  |  |
| --- | --- | --- | --- | --- | --- | --- | --- | --- | --- | --- | --- | --- |
|  |  | KT | NB | KY* | LB | JR | DI | TW | TN | GC | HC* | JO* |
| Aggressor | KT | - | 2 | 0 | 0 | 5 | 1 | 0 | 0 | 3 | 0 | 5 |
|  | NB | 0 | - | 0 | 0 | 7 | 0 | 0 | 0 | 5 | 1 | 6 |
|  | KY* | 0 | 0 | - | 1 | 1 | 0 | 0 | 0 | 0 | 0 | 6 |
|  | LB | 0 | 0 | 0 | - | 0 | 3 | 0 | 0 | 0 | 0 | 0 |
|  | JR | 0 | 4 | 0 | 0 | - | 1 | 0 | 0 | 4 | 0 | 0 |
|  | DI | 0 | 0 | 0 | 0 | 0 | - | 0 | 4 | 1 | 0 | 1 |
|  | TW | 0 | 0 | 0 | 0 | 0 | 0 | - | 0 | 0 | 0 | 0 |
|  | TN | 0 | 0 | 0 | 0 | 0 | 0 | 0 | - | 0 | 1 | 0 |
|  | GC | 0 | 0 | 0 | 0 | 1 | 1 | 1 | 0 | - | 0 | 5 |
|  | HC* | 0 | 0 | 0 | 0 | 0 | 0 | 0 | 1 | 0 | - | 0 |
|  | JO* | 0 | 0 | 1 | 0 | 0 | 0 | 0 | 0 | 2 | 2 | - |

\* Adolescent male

Table 4 shows the numbers of dyadic aggressive interactions expressed and received in each dyad among 10 adult male chimpanzees. Dyadic aggressive interactions were observed in 32 out of 45 dyads. All the individuals except the alpha male were attacked multiple times by higher-ranking males. Of the all 100 aggressive interactions among adult males, 47 were expressed by the alpha male, Goku.

**Table 4.** Aggressive interaction matrix with numbers of aggression expressed and received among 10 adult chimpanzees of the M group.

|  |  | Receiver |  |  |  |  |  |  |  |  |  |
| --- | --- | --- | --- | --- | --- | --- | --- | --- | --- | --- | --- |
|  |  | GK | PO | BR | IC | PR | TK | DO | PT | BL | JO |
| Aggressor | GK | - | 3 | 4 | 9 | 3 | 6 | 5 | 6 | 2 | 9 |
|  | PO | 0 | - | 0 | 0 | 5 | 2 | 0 | 3 | 2 | 2 |
|  | BR | 0 | 0 | - | 0 | 3 | 0 | 0 | 1 | 1 | 0 |
|  | IC | 0 | 0 | 0 | - | 5 | 2 | 1 | 1 | 0 | 2 |
|  | PR | 0 | 0 | 1 | 2 | - | 1 | 2 | 0 | 0 | 0 |
|  | TK | 0 | 0 | 0 | 0 | 1 | - | 3 | 0 | 1 | 1 |
|  | DO | 0 | 0 | 0 | 2 | 1 | 0 | - | 0 | 0 | 0 |
|  | PT | 0 | 0 | 0 | 1 | 0 | 0 | 1 | - | 0 | 0 |
|  | BL | 0 | 1 | 0 | 1 | 2 | 0 | 0 | 1 | - | 1 |
|  | JO | 0 | 0 | 0 | 0 | 0 | 0 | 0 | 0 | 0 | - |

### Comparison of the type and frequency of adult male aggressive interactions between bonobos and chimpanzees

Table 5 shows the number of OHUs observed, the mean number of adult males in the party, the total number of adult males observed in each OHU, and the number of aggressive interactions among adult male bonobos of the E1 group and adult male chimpanzees of the M group (note that this comparison did not include aggressive interactions involving adolescent male bonobos).

**Table 5.** Type and frequency of aggressive interactions among adult males compared between bonobos of the E1 group and chimpanzees of the M group.

| Groups | Total number of OHUs observed | Mean number of males in the party | Total number of males observed | Number of male-male aggressive interactions* |  |  |  |  | Total number of males involved in aggressive interactions | Frequency of aggressive interactions per OHU | Frequency of aggressive interaction per OHU per male |
| --- | --- | --- | --- | --- | --- | --- | --- | --- | --- | --- | --- |
|  |  |  |  | Physical contact | Chasing | Directed display | Others | Total |  |  |  |
| Bonobos | 217 | 3.60 ± 1.80 | 782 | 0 (0) | 25 (14) | 31 (0) | 1 (0) | 57 (14) | 127 | 0.263 | 0.162 |
| Chimpanzees | 677 | 5.67 ± 3.09 | 3840 | 30 (0) | 126 (15) | 42 (9) | 3 (0) | 201 (24) | 429 | 0.297 | 0.112 |
| Chimpanzees (under the presence of females with MS) | 376 | 6.35 ± 2.98 | 2389 | 23 (0) | 99 (13) | 29 (6) | 1 (0) | 152 (19) | 325 | 0.404 | 0.136 |
| Chimpanzees (under the absence of females with MS) | 301 | 4.82 ± 3.03 | 1451 | 7 (0) | 27 (2) | 13 (3) | 2 (0) | 49 (5) | 104 | 0.163 | 0.072 |

Numbers in parentheses represent the number of polyadic aggressive interactions.
\* The 35 agonistic interactions involving adolescent males in bonobos are not included in this table.

Among bonobos, > 50% of aggressive interactions were dyadic directed displays. Chasing behaviors were observed with a similar frequency, but more than half of them were polyadic aggressive interactions (shown in parentheses in Table 5). Aggressive physical contact among adult males was not observed during this study.

Among chimpanzees, chasing was the most frequent aggressive interaction and accounted for approximately 60% of the total aggressive interactions. Aggressive physical contact accounted for 15% of all aggressive interactions, and > 70% of these occurred under the presence of females with MS.

In bonobos, all 14 observed polyadic aggressive interactions among adult males involved chasing behaviors, with multiple individuals pursuing one individual, and at least one adult male participating as an aggressor. Three or four individuals joined as aggressors in eight of the 14 cases, and the aggressee of these interactions was a single male, Jiro in 13 of 14 cases. Kitaro or/and Nobita joined in 12 of these 13 cases of aggression toward Jiro. Females joined as aggressors in six of the 14 cases. Although Kiku joined as an aggressor with Nobita in only one case, probably because she had a newborn baby during this study period, Nao, an old female ally of Kiku, joined in three cases to attack Jiro.

Out of all 24 polyadic aggressive interactions among chimpanzees, 10 cases were coalitionary aggression comprising nine chasing behaviors and one directed aggressive display from two males. All of these coalitionary aggressions occurred under the presence of females with MS, except for one case of chasing behavior. Female chimpanzees did not join any of the coalitionary aggression as aggressors. Chimpanzees also showed 14 aggressive interactions from one male toward multiple individuals. These interactions include six cases of chasing directed at multiple individuals and eight cases of aggressive displays after which other multiple males responded by screaming or fleeing. Such aggressive interactions from one individual to multiple individuals were not observed in bonobos.

The frequency of overall aggressive interactions per OHU in bonobos (0.263) was lower than that of chimpanzees (0.297) but the difference was not statistically significant (chi-square test: χ^2^ = 0.778, df = 1, *p* = 0.378). When data on chimpanzees was categorized into two groups based on the presence/absence of females with MS, the frequency of aggressive interactions followed the order: chimpanzees under the presence of females MS (0.404) > bonobos (0.263) > chimpanzees under the absence of females with MS (0.163). These differences were statistically significant (chi-square test: χ^2^ = 48.41, df = 2, *p* < 0.001). The results of the Bonferroni-corrected *post hoc* tests showed that the differences were significant in all three pairs: bonobos vs. chimpanzees with females with MS (*p* < 0.01), bonobos vs. chimpanzees without females with MS (*p* < 0. 001), chimpanzees with females with MS vs. chimpanzees without females with MS (*p* < 0.001) (Table 6).

**Table 6.** The results of the Bonferroni-corrected post hoc tests.

| Group comparison | Adjusted p-value |  |
| --- | --- | --- |
|  | Aggression per OHU | Aggression per OHU per male |
| Bonobos vs. Chimpanzees | NS | <0.05 |
| Bonobos vs. Chimpanzees (under the presence of females with MS) | <0.01 | NS |
| Bonobos vs. Chimpanzees (under the absence of females with MS) | <0.001 | <0.001 |
| Chimpanzees (under the presence of females with MS) vs. Chimpanzees (under the absence of females with MS) | <0.001 | <0.001 |

However, when the difference in the mean number of males in the party was considered, these differences between the species/conditions changed. The frequency of aggressive interactions per male in bonobos (0.162) was higher than that of chimpanzees (0.112), and the difference between the species was statistically significant (chi-square test: χ^2^ = 4.821, df = 1, *p* = 0.028). It seemed to be because a larger number of males attended OHU in chimpanzees, but aggressive interactions specifically occurred among some higher-ranking males, lowering the per-male frequency of aggressive interactions in chimpanzees. When data on chimpanzees were categorized based on the presence/absence of females with MS, the relationships in the frequencies of aggressive interactions per male was in the order: bonobos (0.162) > chimpanzees under the presence of females with MS (0.136) > chimpanzees under the absence of females with MS (0.072), and the differences were statistically significant (chi-square test: χ^2^ = 20.471, df = 2, *p* < 0.001). The results of the Bonferroni-corrected *post hoc* tests showed that the differences were significant in bonobos vs. chimpanzees under the absence of females with MS (*p* < 0. 001) and chimpanzees under the presence of females with MS vs. chimpanzees under the absence of them (*p* < 0.001), but not in bonobos vs. chimpanzees under the presence of females with MS (*p* = 0.410) (Table 6).

## Discussion

In this study, we have focused on the difference in the frequency and intensity of male aggression between chimpanzees and bonobos. We hypothesized that the patterns of association and behaviors of male bonobos are strongly affected by the prolonged sexual receptivity of females and the constant presence of multiple receptive females (females with MS) in the group (Kano 1992; Furuichi and Hashimoto 2002; Hashimoto et al. 2022; Furuichi 2023), and the long-lasting mother–son relationships and strong support from mothers for mating competition and acquisition of the alpha status (Furuichi 1989; Kano 1992; Surbeck et al. 2010; Furuichi 2011; Ishizuka et al. 2018; Furuichi 2019; Surbeck et al. 2019; Shibata and Furuichi 2023). We examined the four predictions derived from this hypothesis.

The results of the current study support the first prediction that male bonobos find a higher number of receptive females in their parties than male chimpanzees do. Chimpanzee and bonobos males found 0.47 ± 0.68 and 1.22 ± 1.15 receptive females in each OHU, respectively, probably because a large proportion of females in the group are receptive showing MS due to the higher proportion of the receptive period of females in bonobos, especially caused by the resumption of MS in the early stage of postpartum amenorrhea (Furuichi and Hashimoto 2002; Hashimoto et al. 2022).

While at least one female with MS was found in 94.6% of all observation days in bonobos, they were found in only 55.4 % of observation days in chimpanzees. On the days when females with MS were not observed in the group, the modal value of the number of male chimpanzees in parties was one, indicating that the males were highly dispersed. In chimpanzees, low-ranking males were substantially more likely to range alone or in parties without other males than high- and middle-ranking males in the absence of receptive females, which might help them to reduce the probability of being attacked by other males (Shibata et al. 2022). However, on the days when females with MS were observed, nearly half of the male chimpanzees were observed in the same party, similar to male bonobos in pooled data. In such days when females with MS were observed in the group, no significant relationship between solitary ranging and male rank was observed in chimpanzees, suggesting active participation by even low-ranking males in the party (Shibata et al. 2022). Despite a significant difference in the distribution of the number of adult males in the party between bonobos and chimpanzees, both species seem to show a similar tendency to attend mixed-sex parties when females with MS were present. This result underscores the significant impact of the presence (in chimpanzees) or absence (in bonobos) of days without females with MS on the social dynamics among males.

The analysis of aggressive interactions among adult males did not fully support the second prediction, which posited that the frequency of aggressive interactions is lower in bonobos than in chimpanzees (Tables 5 and 6). In particular, while the frequency of aggressive interactions per OHU showed no significant difference between bonobos and chimpanzees, the frequency of aggressive interactions per OHU per male was significantly higher in bonobos than in chimpanzees. When comparing the frequency of aggressive interactions in bonobos with those in chimpanzees under the presence of females with MS, the frequency per OHU was significantly lower in bonobos than in chimpanzees but there was no significant difference in the frequency per OHU per male (Tables 5 and 6). In the days when females with MS were not observed in chimpanzees, both the frequency of aggressive interactions per OHU and per OHU per male were significantly lower than those in pooled data of bonobos and those in chimpanzees under the presence of females with MS (Tables 5 and 6), probably because aggressive interactions did not occur over access to MS females, and low- ranking males frequently range alone or in parties without other males to avoid aggression of higher-ranking males under such circumstances (Shibata et al. 2022).

Conversely, prediction 2 was supported when examining the type of aggression. In bonobo males, aggressive interactions were primarily characterized by chasing or aggressive displays, and notably, unlike in chimpanzees, no instance of aggressive physical contact was observed during this study. This may explain why lower-ranking bonobo males do not need to leave the party to avoid attacks by dominant males. Although a single male was targeted in all polyadic aggressive interactions in bonobos, one male targeted multiple males in 14 of 24 cases of polyadic aggressive interactions in chimpanzees. These interactions occurred while other males were resting or grooming in proximity, and the alpha male was the aggressor in nine of these cases. This behavior aligns with what has been previously termed as “separating intervention,” in other studies, wherein alpha males prevent other males from forming affiliative relations and coalitions (de Waal 1982; Nishida and Hosaka 1996). This underscores the intricacies of male competition among chimpanzees and the substantial impact of aggressive behaviors from dominant individuals.

The results of this study support the third prediction, indicating that aggressive interactions between males over access to receptive females are less frequent in bonobos than in chimpanzees. Previous research on chimpanzees has documented an increase in male aggression in the presence of receptive females (Muller and Wrangham 2004). Similarly, in the current study, we observed a rise in the frequency of more severe aggressive interactions, such as aggressive physical contact and chasing behavior, among chimpanzees when females with MS were present. Notably, nine of 10 cases of coalitionary aggressive interactions in chimpanzees occurred under the presence of females with MS. These results suggest that aggressive interactions among male chimpanzees intensify under the presence of receptive females, and males strategically form coalitionary attacks for mating competition. In contrast, a previous study at Wamba reported a relatively low frequency of aggressive behavior that interferes with the mating of subordinates in bonobos (27 of 515 cases of copulations (Kano 1996); 2 of 36 cases of agonistic interactions (Furuichi 1997)). Similarly, in this study, out of 73 cases of aggressive interactions in clear contexts, only seven cases were observed over access to females with MS (Shibata, unpublished data).

The findings of the current study also corroborate the fourth prediction, indicating that aggressive interactions between males involve their mothers more frequently in bonobos than in chimpanzees. In chimpanzees, aggressive interactions occurred in 32 out of 45 dyads of adult males, with all males except the alpha male being aggressed multiple times by higher-ranking males. However, females did not participate in any of the aggressive interactions among males. On the contrary, in bonobos, dyadic aggressive interactions were observed in only 22 out of 55 dyads of adult and adolescent males. Notably, dyadic aggression was directed toward two specific males in as many as 37 out of 76 cases. During this study period, the two sons of the second-ranking female Jacky (Jo and Jiro) frequently challenged the sons of Kiku (Kitaro, Nobita, and Kiyota) and faced counter-attacks. Jiro, in particular, was targeted in 13 out of 14 polyadic aggressive interactions, and Kitaro or/and Nobita joined in 12 of these 13 cases of aggression toward Jiro, highlighting the intensity of competition for the alpha male position. These observations suggest that a significant proportion of aggressive interactions among bonobo males involved competition for the alpha male position. In all 45 cases of aggressive interactions between the sons of Kiku and the sons of Jacky, Kiku or her female ally, Nao, joined in four cases to attack Jiro. In the PE group neighboring the E1 group, the mother of one of the males participated in approximately 6% of aggressive interactions between males (Tokuyama, personal communication). Another study at LuiKotale reported that mothers joined approximately 22% of aggressive interactions over access to receptive females (Surbeck et al. 2010). The current study, along with these preceding reports, indicates that bonobo females participate in aggressive interactions more frequently than female chimpanzees do, intervening in mating competitions or competitions for alpha-male status involving their sons.

It should be noted that although a part of the second prediction was not supported due to the equal or even higher frequency of aggressive interactions among male bonobos compared to male chimpanzees, the frequency of aggressive interactions among male bonobos substantially decreases when interactions among these specific males are excluded.

In conclusion, the scrutiny of the four predictions aligns with our hypothesis that the prolonged sexual receptivity of females exert a significant influence on male agonistic relationships in bonobos. As documented in previous studies, the operational sex ratio (proportion of adult males to sexually receptive females) in bonobos is notably reduced due to the early resumption of sexual swelling and receptivity during the infertile nursing period (Kano 1992; Furuichi and Hashimoto 2002; Hashimoto et al. 2022; Furuichi 2023). This study reveals that this reduction increases the presence of females with MS in the group, along with the number of such females that male bonobos encounter in the parties they attend. This heightened opportunity for mating appears to contribute to less frequent aggressive interactions, particularly those related to access to receptive females, and a decrease in the severity of aggressive behaviors among males.

Additionally, our findings highlight that a significant portion of aggressive interactions among male bonobos revolves around struggles for the alpha-male position and that female bonobos sometimes attend to aggressive interactions to support their sons. The alpha female, Kiku, was observed to participate in these interactions only once during this study, likely influenced by her caring for a newborn infant. However, Kiku’s female ally, Nao, consistently joined in aggressive interactions to support Kiku’s sons. Examining the historical alpha status dynamics in the E1 group over nearly the past 40 years (1983–2020) emphasizes the substantial impact of the mother’s status among females and their intervention in struggles for alpha status among male bonobos (Furuichi 1997, 2011; Shibata and Furuichi 2023). This seems to be a reason why relatively young males (late adolescent or young adult) with mothers in their prime age tend to acquire the alpha male status in bonobos (Furuichi 1997, 2019; Shibata and Furuichi 2023) In contrast, in chimpanzees, age emerges as a pivotal factor in acquiring alpha status among males. Hasegawa and Kutsukake (2015), based on data from three long- term study sites, reported that male chimpanzees typically reach their highest potential to become alpha at an age of approximately 25 years, except in cases where the alpha status is achieved through coalition formation. Eastern chimpanzees in Mahale acquire alpha status at the age of 19 to 37 years (Nishida 2008). In Gombe, middle-aged males reach their highest rank, dominating most, if not all, younger and older males. While cases have been reported of males with supportive elder brothers becoming alpha, and mothers occasionally support their offspring in fights, the influence of such maternal support on the rank of adult males remains unclear (Bygott 1979).

The variations in the impact of maternal support observed between the two species may stem from differences in mother–son relationships. In chimpanzees, males significantly decrease their association with mothers when mothers resume cyclic receptivity, eventually severing their relationship to associate with male adults (Pusey 1983; Hayaki 1988). Conversely, male bonobos maintain close associations with their mothers into adulthood, remaining in the same party, providing ample opportunities for maternal support (Furuichi 1989; Kano 1992; Surbeck et al. 2010; Schubert et al. 2013; Hashimoto and Furuichi 2015). However, it remains unclear whether the prolonged mother–son associations enhance the influence of mothers on intermale competition or if the evolution of female behaviors to support their sons leads to such enduring mother–son associations.

As highlighted in the Introduction, a question in male reproductive competition is the more pronounced reproductive skew toward high-ranking males in bonobos compared to chimpanzees, despite seemingly less despotic intermale relationships (Surbeck et al. 2017b; Ishizuka et al. 2018; Surbeck et al. 2019). Particularly in the E1 group, 10 out of 11 infants with confirmed paternity were sired by alpha males at that time, with nine cases by Nobita and one by Kitaro, both sons of the alpha female Kiku (Ishizuka et al. 2018; Ishizuka 2023). As explained above, one of the reasons for the high reproductive skew in bonobos might be the direct support of high-ranking mothers in the mating competition to support their high-ranking sons. Although aggressive interactions among males do not occur frequently, the presence of high-ranking mothers or their aggressive intervention may suppress the mating behaviors of their sons’ rivals (Surbeck et al. 2010; Furuichi 2023). Another reason might be the characteristic spatial distribution of bonobos within the mixed-sex party. Unlike female chimpanzees, female bonobos gather and stay at the center of the party, and males tend to remain peripheral (Kuroda 1979; Furuichi 1989; Hashimoto and Furuichi 2015). However, due to the strong mother-son association, sons of high-ranking females tend to stay in the gathering of females with their mothers and have more opportunities or easier access to receptive females even if they do not suppress mating behaviors of lower-ranking males by aggressive interactions (Surbeck et al. 2012; Yokoyama and Furuichi 2022; Furuichi 2023). Although these possibilities are yet to be confirmed in future studies, such high influences of support by high-ranking females seem to diminish the significance of direct aggressive interactions among males and reduce the severity of aggressive behaviors. Additionally, female behaviors for supporting their sons might have evolved in bonobos for their own reproductive success, aiming to increase the number of their grandchildren (Furuichi 1997; Surbeck et al. 2019; Furuichi 2023).

The present study faces a limitation stemming from the truncated observation period for bonobos, a consequence of the COVID-19 pandemic. In contrast to the comprehensive eight-month study duration for chimpanzees covering a majority of the year, the investigation into bonobos spanned a condensed four-month interval, specifically from September to December. This temporal discrepancy raises concerns, particularly if male behavioral patterns are influenced by seasonality, particularly in relation to fruit abundance. Studies on a neighboring group at Wamba, PE, indicated that intergroup encounters were most frequent during the early peak of fruit abundance, coinciding with a reduction in aggression between males within the group (Sakamaki et al. 2018; Tokuyama et al. 2019). Conversely, a study at Kokolopori, a nearby site to Wamba, reported an increase in aggression and cortisol levels in males with rising fruit abundance (Cheng et al. 2021). Fortunately, the four-month period of our study aligned with the transition from the highest to the lowest fruiting season at Wamba (Mulavwa et al. 2008); therefore, the current results may not significantly biased by the seasonal change in fruit abundance. However, through studies with with extended periods, we must assess the influences of seasonality in the abundance and distribution of food resources on the type and frequency of intermale aggressive interactions.

It is crucial also to note that the behavioral tendencies observed in this study cannot be universally generalized across populations of bonobos and chimpanzees. Intermale relationships are likely influenced by various factors beyond fruiting seasonality, including group size (Mitani et al. 2000; Watts 2018), the personalities of high-ranking males and females (Goodall 1986; Takahata 1990; Shibata and Furuichi 2023), and differences between chimpanzee subspecies (Boesch 1996). Notably, intermale relationships in bonobos at Wamba vary across years depending on the presence or absence of competing mother–son pairs and their respective personalities. Hence, additional studies involving multiple groups of bonobos and chimpanzees across various years are essential for a comprehensive understanding of behavioral tendencies common to each species and the variations in intermale relationships within male- philopatric societies of the *Pan* species.

## Acknowledgments

We thank the National Forestry Authority of Uganda, the Uganda National Council for Science and Technology, and the Ministry of Scientific Research of the Democratic Republic of the Congo for their permission to conduct this research. We thank the local assistants for their help with field observations. We thank Nahoko Tokuyama, Tetsuya Sakamaki, Kazuya Toda, Chie Hashimoto, Hiroyuki Takemoto, Daisuke Shimizu, Akito Toge, and Shintaro Ishizuka for their help during the field research and their valuable discussions. This research was financially supported by the Leading Program in Primatology and Wildlife Science of the Kyoto University, the Japan Society for the Promotion of Science (JSPS) Grant-in-Aid for JSPS Fellows, and JSPS Grants-in-Aid for Scientific Research (to TF, 16H02753, 25304019, and 26257408). We also appreciate the anonymous reviewers for their helpful comments on this paper.

## Author contributions

SS designed this study, collected and analyzed behavioral data, and wrote and revised the manuscript. TF supervised SS in designing this study. Both authors revised the manuscript and gave the final approval for publication.

## Ethical approval

This study was approved by the National Forestry Authority of Uganda and the Uganda National Council for Science and Technology.

## Notes

### Competing Interest Statement

The authors have declared no competing interest.

### Summary of Updates

The manuscript has been updated through the reviewing process of the journal.

